# A Machine Learning Approach to Predicting Dyspnea with Noninvasive Biomarkers

**DOI:** 10.1101/2025.08.15.669955

**Authors:** Karapet G. Mkrtchyan, Anser Qazi, Borena Lonh, Andrew Dong, Gustavo O Ramirez, Mona Eskandari, Shujie Ma, Wei Vivian Li, Erica C. Heinrich

**Affiliations:** Division of Biomedical Sciences, School of Medicine, University of California, Riverside, Riverside, CA, USA; Department of Mechanical Engineering, University of California, Riverside, CA, USA; Department of Statistics, University of California, Riverside, Riverside, CA, USA

**Keywords:** dyspnea, monitoring, critical care, hypoxia, hypercapnia

## Abstract

**Rationale:** Dyspnea is the subjective sensation of breathing discomfort. This symptom is highly prevalent in patients with chronic and critical illness, and its presence is associated with poor clinical outcomes and long-term psychological trauma. The multidimensional nature of the neurophysiological mechanisms underlying dyspnea, paired with individual variation in its presentation, makes identifying and monitoring this symptom difficult, particularly in non-communicative patients.

**Objective:** Undetected and untreated dyspnea in critically ill patients is a significant problem contributing to patient suffering. Therefore, the objective of this study was to investigate the feasibility of machine learning methods for assessing and continuously monitoring dyspnea using easily obtained noninvasive biomarkers.

**Methods:** We recruited healthy participants (N = 60, 35 women) and stimulated dyspnea using a forced end-tidal semi-rebreathing circuit to modulate arterial oxygen and carbon dioxide levels while collecting non-invasive biomarker data and continuous self-reported dyspnea severity scores. This data was used to train machine-learning models to predict the presence or absence of significant dyspnea (Numeric Rating Scale ≥ 3). We then compared the performance of our final model to observational estimates by trained healthcare providers.

**Measurements and Main Results:** The final model (Random Forest) performed well (PR-AUC=0.832) and exceeded the accuracy of observation estimates made on the same participants using the Respiratory Distress Observational Scale (RDOS) (accuracy=54%).

**Conclusions:** These results indicate that machine learning models can utilize non-invasive biomarker inputs to accurately predict carbon dioxide- and hypoxia-induced dyspnea in a healthy population during spontaneous breathing.

## INTRODUCTION

Dyspnea is the subjective experience of breathing discomfort and, when left untreated, can produce profound patient suffering.^1–4^ This symptom is highly prevalent in patients with both acute and chronic respiratory and cardiovascular disease and is an independent predictor of mortality in these groups.^5,6–9^ Dyspnea causes delays in extubation in mechanically ventilated patients, and is correlated with an increase in noninvasive ventilation failure, generally leading to worse outcomes.^10^ It can also cause lingering damage to psychological health well after physiological symptoms have subsided, and hospitalized patients who experience severe dyspnea are at high risk of developing depression, anxiety, or PTSD-like symptoms after their hospitalization.^1,11–14^

In the past decades, much progress was made in our understanding of the pathophysiology of dyspnea. Several qualitatively distinct sensations contribute to this symptom, including work of breathing, chest tightness or bronchoconstriction, and “air hunger” or unsatisfied inspiration.^6,15,16^ These sensations are driven by several neural inputs which modulate respiratory drive, including information from chemoreceptor stimulation, chest wall mechanoreceptors, and vagal C fibers.^16–20^ Dyspnea occurs when there is a mismatch between the expected result of central neural output and the actual perceived airflow or ventilation.^6,17^

Despite the widespread presence of dyspnea and its significant impact on patient quality of life, the detection of dyspnea in the hospital setting continues to be challenging. Current methods of detection, such as the respiratory distress observation scale (RDOS), rely heavily on subjective measures (such as look of fear) and observations that can be masked due to patient condition or medications administered (such as grunting and nasal flaring).^21^ Furthermore, healthcare providers often “woefully” underestimate dyspnea severity based on observation alone.^15,22–25^ In many cases, patients may be unable to communicate their discomfort, leading to unrecognized patient suffering.^4^ It is likely that this disparity is due to the complex, multivariate nature of dyspnea, as well as individual variation in how patients experience, display, and describe their discomfort.

There is an urgent need for a clinical tool to improve detection of dyspnea and continuously monitor this symptom. Such a tool would empower physicians with novel information about patient discomfort and indicate when interventions are needed. To address this need, we aimed to determine if self-reported dyspnea severity can be accurately predicted using noninvasive biomarkers. To accomplish this, we experimentally stimulated dyspnea while measuring several easily monitored physiological factors and used a machine learning approach to create a model to predict self-reported dyspnea severity using these biomarkers as inputs. We hypothesized that self-reported dyspnea could be accurately predicted with these simple biomarkers and that this model would score higher on performance metrics such as precision and recall than healthcare providers’ observational estimates.

Some of the results of these studies have been previously reported in the form of abstracts.^26,27^

## METHODS

### Participants

This study was approved by the University of California, Riverside Clinical Institutional Review Board (IRB #22063) and was conducted in accordance with the *Declaration of Helsinki*, except for registration in a database. 60 healthy individuals were recruited (35 women, 25 men, 23.10 ± 5.83 years of age). Full inclusion criteria are provided in the supplemental methods.

### Dyspnea experiments

Dyspnea was stimulated by exposing participants to six different combinations of carbon dioxide and oxygen (**Table1**) using a partial-rebreathing end-tidal forcing system as described previously.^28^ Trials were administered in a randomized, participant-blinded order, each lasting four minutes and interspaced by four-minute rest periods where the participants breathed room air (**Figure 1**).

**Figure 1.**
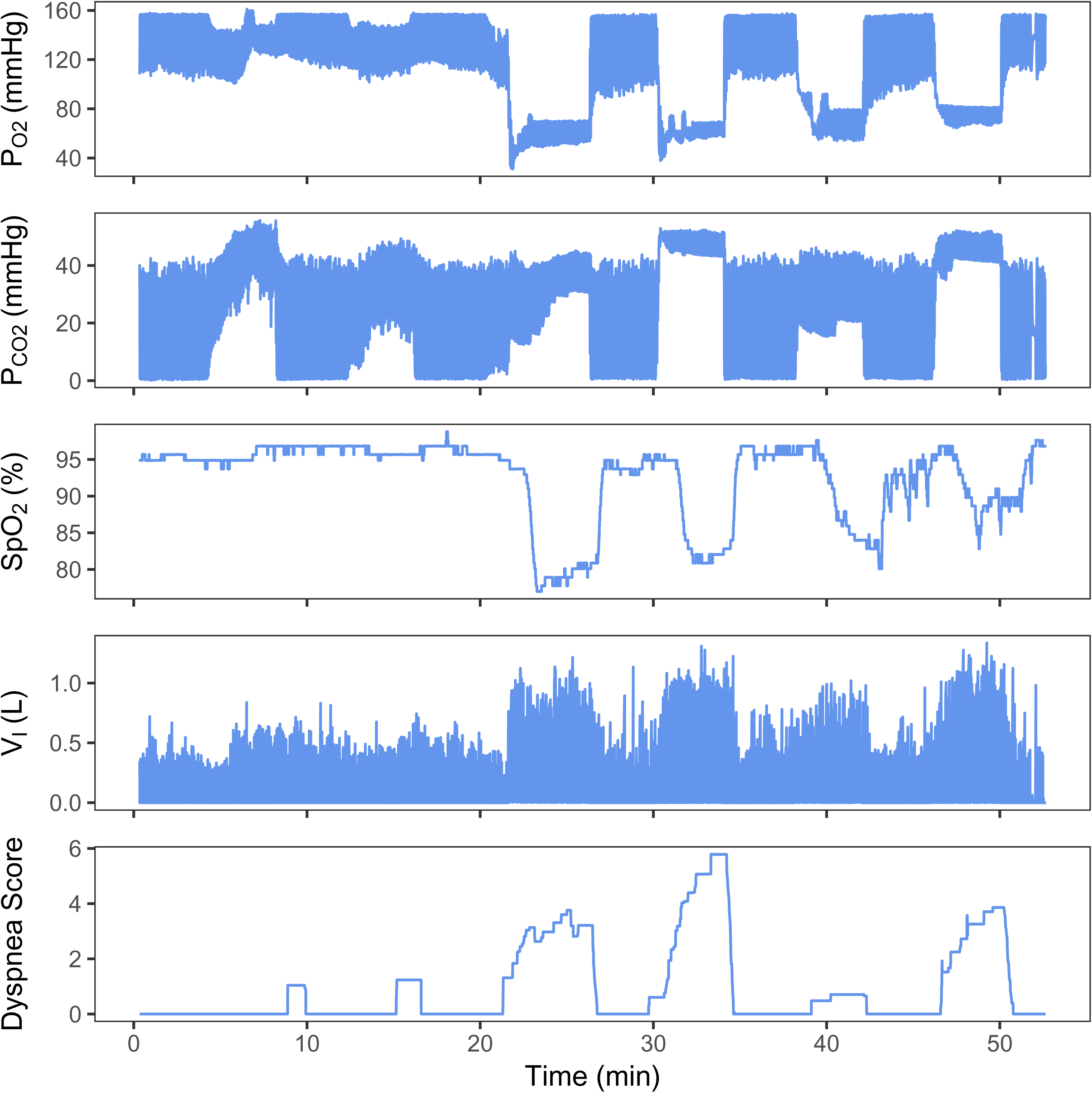
Experimental treatments. A representative raw data trace of a complete experimental procedure in one participant. P_O2_ and P_CO2_ levels are measured directly in front of the mouth. V_I_ peaks represent inspirations. Dyspnea scores are self-reported NRS scores recorded continuously throughout the test.

**Table 1.** Treatment targets for end-tidal forcing experiments. Treatments were administered in a randomized, participant-blinded order during experiments.

| **Trial** | **Inspiratory**<br>**P<sub>O2</sub> (mmHg)** | **End-tidal**<br>**P<sub>CO2</sub> (mmHg)** |
| --- | --- | --- |
| 1 | 149 | 45 |
| 2 | 149 | 52 |
| 3 | 80 | 45 |
| 4 | 80 | 52 |
| 5 | 68 | 45 |
| 6 | 68 | 52 |

SpO_2_, ECG, partial pressures of oxygen (pO_2_) and carbon dioxide (pCO_2_), and respiratory flow were measured continuously. Self-reported dyspnea severity was provided by the participants on a 0-10 scale (NRS) either at the end of the four-minute trial or continuously using a knob to modify the score on a screen in front of them. Biomarker and dyspnea score data was extracted in one-minute increments at the end of each four-minute trial and rest period.

### Physician dyspnea severity rating

Participants were video recorded from a front-facing and profile view during the experiments. 30-second clips were collected and paired with simultaneous screen recordings of heart and respiratory rates. A custom MATLAB program was created to display the clips one-by-one and prompt the viewer to score criteria of the RDOS. MICU and ED physicians and nurses completed these ratings (n =19). Each video was labeled with “presence of dyspnea” if the total RDOS score was equal to or above the threshold used. Initial threshold was set to 3. Due to difficulty observing nasal flaring, this criterion was removed.

### Statistical analysis and machine learning approach

All algorithms were developed using the Anaconda distribution of packages for python, models were carried out using the Sci-kit Learn toolkit, and models were re-validated using the same parameters in R. Data was quality checked, pre-processed and coded, and missing data was removed listwise. Models included dyspnea severity as the supervised label and input variables including sex, BMI, age, and mean, maximum, minimum, and standard deviation of the following: end tidal pCO_2_, inspiratory pO_2_, heart rate, total ventilation, tidal volume, respiratory rate, inspiratory flow rate, and SpO_2_. Dyspnea severity was divided into two categories (present or absent), with a variety of thresholds tested for optimal performance. Six classification models were chosen to evaluate baseline for model performance. Cross-validation was done with a 30-70 split of the data for testing and training, respectively. Random forest was selected as the top-performing model to evaluate based on results.

## RESULTS

### Dyspnea severity as a function of gas treatments

Self-reported dyspnea severity differed significantly across the 6 experimental treatments (main effect: F(6,582)=136.5, p<0.001) (**Figure 2**). These differences were driven by significant main effects of both inspired pO_2_ (F(2,584)=47.4, p<0.001) and end-tidal pCO_2_ (F(2,584)=142.7, p<0.001). There was substantial variation in self-reported dyspnea severity across participants even under the same experimental treatments (**Figure 2**, **Table S1**). This is not driven by variance in the achieved pO_2_ and pCO_2_ treatment levels using the end-tidal forcing system. We achieved our target gas tensions within an average standard deviation inter-trial of ± 1.1 mmHg and intra-trial of ± 0.6 mmHg for end-tidal pCO_2_, and inter-trial of ± 3.0 mmHg for and intra-trial of ± 2.6 mmHg for inspired pO_2_ (**Figure S1**). Actual end-tidal pCO_2_ and pO_2_ values were utilized in machine learning models rather than grouping by treatment.

**Figure 2.**
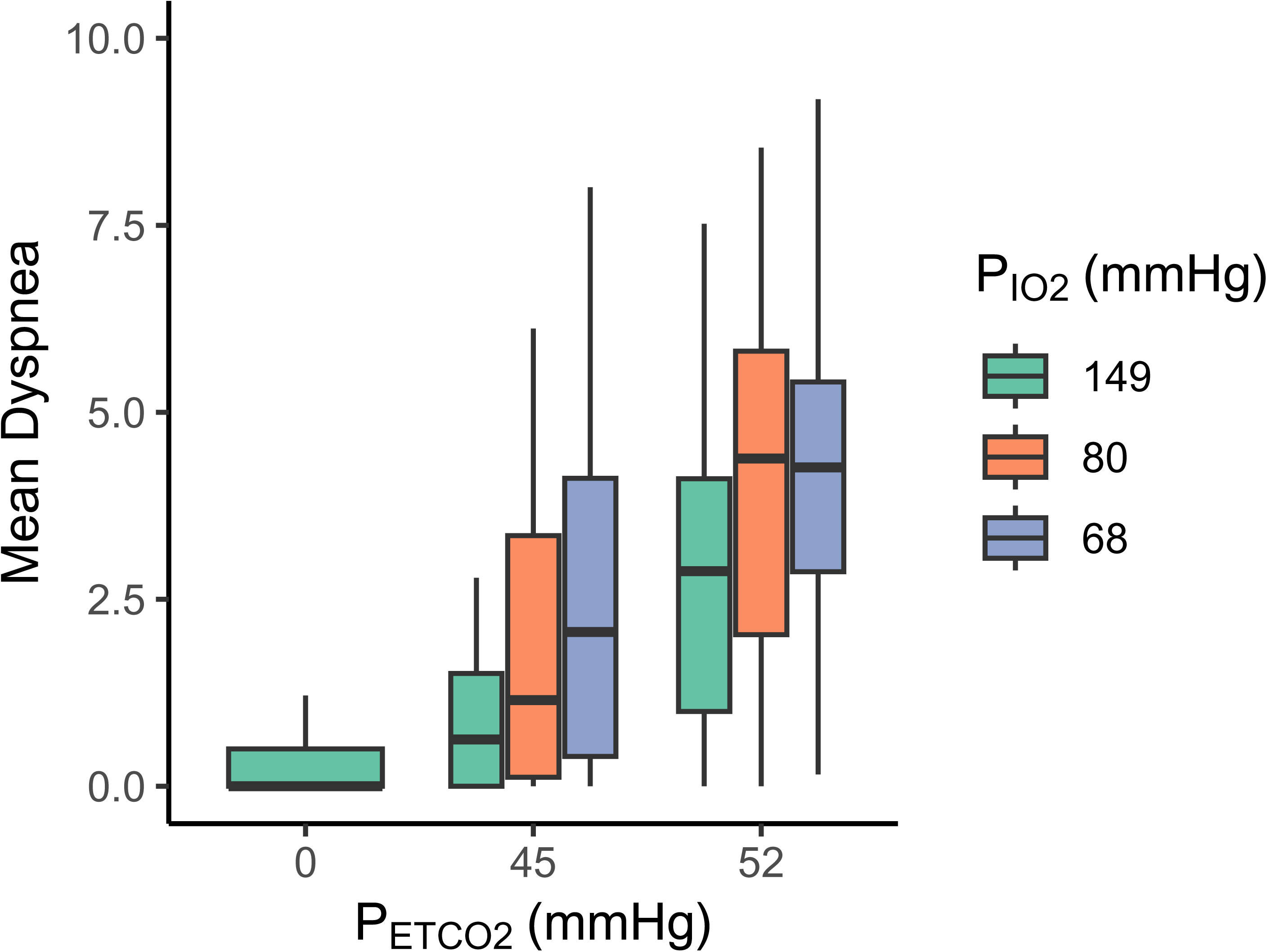
Self-reported dyspnea severity as a function of gas treatments. Mean self-reported dyspnea severity (via NRS) increased as a function of end-tidal P_CO2_ and inspired P_O2_. Tukey-style boxplots are provided with means indicated by thick center bars, first and third quartiles represented by boxes and whiskers representing 1.5 * IQR. Outlier points are not plotted.

### Efficacy of traditional individual biomarkers

**Table 2** provides a summary of the independent effects of key biomarkers on mean dyspnea severity under each treatment via a mixed linear model which considers age, sex, BMI, end-tidal pCO_2_ and inspiratory pO_2_ as covariates. All biomarkers except SpO_2_ show significant main effects.

**Table 2.** Summary of main effects of individual biomarkers in mixed linear models including sex, BMI, age, and gas tensions as covariates. P values represent the significance level of the main effect of the individual biomarker in the model.

| Biomarker | $\beta$ | P | Std. Error | CI (0.025%) | CI (0.975%) |
| --- | --- | --- | --- | --- | --- |
| Respiratory rate (breaths/min) | 0.102 | <0.001 | 0.018 | 0.066 | 0.139 |
| Tidal Volume (L) | 1.682 | <0.001 | 0.263 | 1.166 | 2.197 |
| Minute Ventilation (L/min) | 0.140 | <0.001 | 0.014 | 0.114 | 0.167 |
| Heart rate (beats/min) | 0.028 | 0.006 | 0.010 | 0.008 | 0.048 |
| SpO<sub>2</sub> (%) | 0.036 | 0.158 | 0.026 | -0.014 | 0.087 |

### Selection of cut-off for binary classification of dyspnea during model training

NRS scores of two through five were tested separately as cut-off points for classification due to the variety of cut-off points utilized in prior studies, though cut-offs of three to four are generally recommended for distinguishing moderate discomfort from mild/lack of.^29,30^ A cut-off of three was ultimately selected based on highest performance. **Table S2** summarizes model performance across the various thresholds. PCA analysis shown in **Figure S2** also indicates more clear division of groupings with a dyspnea score greater than or equal to three.

### Categorical prediction models

All models performed with high accuracy; **Figure3A** shows the testing results of all models. Area under the receiver operating characteristic curve (ROC-AUC) values were highest for the KNN (0.926) and random forest model (0.923) with all models performing with ROC-AUC values above 0.85. Due to the imbalanced nature of the results (presence of dyspnea severity ≥3 was less common), area under the precision-recall curve (PR-AUC) values were also calculated (**Figure 3B**). Random forest performed the best with an PR-AUC of 0.832, with the nearest competitor being SVM at 0.809. Since the random forest model demonstrated highest performance, it was chosen as the focus of subsequent analyses. **Figure 3C** provides feature importance scores for the top ten biomarkers within the final random forest model. Standard deviation of flow rate, mean total ventilation and mean inspiratory flow rate were the top three features in this model. Beyond the top three features there is an initial sharp decline in feature importance, and the remaining features in the top 10 exhibit similar levels of importance.

**Figure 3.**
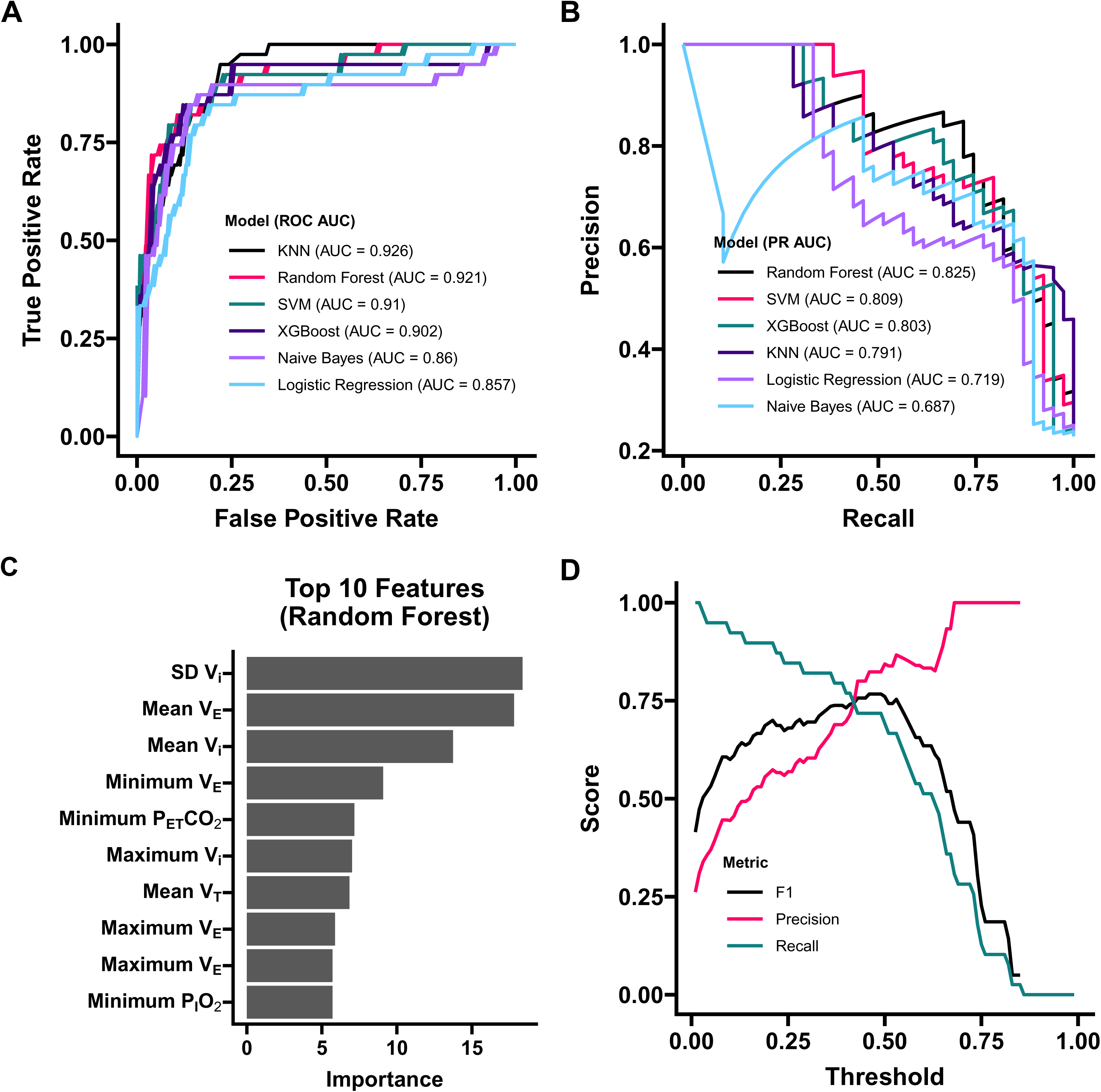
Model testing performance and metrics. **(A)** Receiver Operating Characteristic of all models. **(B)** Precision-Recall curves **(C)** Feature Importance of the top 10 features of the random forest model. **(D)** Precision and Recall trade-off shown with corresponding F1 scores for random forest model. Threshold of 0.42 determined as ideal based on model performance.

In a separate analysis, we sought to verify if the final random forest model’s performance was biased across sex groups. To do this, we tested the final model separately on test datasets derived only from male or female participants. We found no significant difference in performance for the prediction between sex groups (Male: precision = 0.980, recall = 0.909, F1 = 0.943; Female: precision = 0.915, recall = 0.974, F1 = 0.943; p = 1).

Next, we determined the threshold on the predicted probabilities for classification (**Figure 3D**). A threshold of 0.42 lead to the highest F1 score (0.805). Identifying patients who have dyspnea (positive result) is of key importance, so further experiments and clinical verification is needed to determine the ideal threshold balance between precision and recall in this application. For example, recall can be improved if the classification threshold is further reduced, at the expense of precision, which may be of value in an ICU setting where false negative estimates are harmful.

### Physician and nurse performance

Physicians and nurses performing RDOS scoring on participants from the machine learning dataset achieved an accuracy of 58.82% using standard thresholds. The standard threshold of ≥3 was used for the RDOS, but a variety of self-report cut-offs were tested.^31^ **Table 3** shows the confusion matrix and overall physician performance. Notably, recall rate was alarmingly low and was the primary reason for loss in accuracy. Recall performance did not improve across a variety of self-reported dyspnea score cut offs, although precision did improve markedly as the cut off was decreased. F1 and accuracy scores remain largely unchanged across all cut off points. As literature has expanded in recent years on the use of the RDOS and it has been adapted to various applications with changes in scale and cut off points, we also tested performance across a variety of RDOS thresholds.^32–34^ **Table S3** shows the complete set of performance results. Generally, decreases in the RDOS threshold greatly increase recall with minimal loss of precision.

**Table 3.** Healthcare provider performance in identifying the presence of significant dyspnea, using standard RDOS threshold of ≥3 for the observational estimate of presence of dyspnea compared at several different NRS levels of self-reported dyspnea severity.

| **NRS**<br>**Dyspnea**<br>**Cut-Off** | **True**<br>**Pos.** | **True**<br>**Neg.** | **False**<br>**Pos.** | **False**<br>**Neg.** | **Precision** | **Recall** | **F1** | **Accuracy** |
| --- | --- | --- | --- | --- | --- | --- | --- | --- |
| **2** | 27 | 11 | 2 | 28 | 0.931 | 0.491 | 0.643 | 0.559 |
| **2.5** | 24 | 13 | 5 | 26 | 0.828 | 0.480 | 0.608 | 0.544 |
| **3** | 24 | 13 | 5 | 26 | 0.828 | 0.480 | 0.608 | 0.544 |
| **3.5** | 22 | 18 | 7 | 21 | 0.759 | 0.512 | 0.611 | 0.588 |
| **4** | 22 | 18 | 7 | 21 | 0.759 | 0.512 | 0.611 | 0.588 |
| **4.5** | 17 | 22 | 12 | 17 | 0.5862 | 0.5 | 0.540 | 0.574 |
| **5** | 17 | 22 | 12 | 17 | 0.5862 | 0.5 | 0.540 | 0.574 |

## DISCUSSION

### Individual variation in hypercapnia and hypoxia-induced dyspnea during free breathing

Our data demonstrated that both end-tidal pCO_2_ and inspired pO_2_ levels were associated with self-reported dyspnea severity in free-breathing healthy individuals (**Figure 2**). These data support prior findings that hypoxia and hypercapnia induce dyspnea.^16,18,35,36^ While our experimental treatments were effective in stimulating significant dyspnea in this population, we note that there was substantial variation in severity across participants under the same treatment conditions. In contrast, self-reported dyspnea severity is more severe and less variable across individuals in other studies utilizing fixed ventilation protocols.^37^ This may be due to several factors, including feedback from lung mechanoreceptors during larger tidal volumes in voluntary spontaneous breathing, alleviating some of the dyspnea stimulus. Based on these differences in dyspnea severity in free breathing individuals versus those experiencing fixed tidal volumes and/or frequency, future work will be required to validate our models in clinical populations to facilitate translation.

### Predictive power of single biomarkers

Our findings are consistent with prior work showing significant relationships between select individual biomarkers and dyspnea severity. Tachypneic breathing is often viewed as one of the main indicators of dyspnea, and is one of the scored criteria on the RDOS showing the highest predictive value.^38^ This link is clear in previous experiments by Izumizaki et al. (2011) who show that the threshold for dyspnea is positively associated with the threshold for increases in respiratory rate during CO_2_ rebreathing.^39^ Additionally, heart rate shows modest correlations with dyspnea severity in both Campbell’s original RDOS work and our current study.^38^ Tidal volume and minute ventilation were included as biomarkers in our study but are not included in the RDOS. Interestingly, tidal volume had the largest predictive value in our study, which supports work by many others demonstrating the large independent impact of tidal volume on dyspnea in ventilated patients, with low-tidal volume and permissive hypercapnia often favored to prevent lung injury but leading to increased patient discomfort.^40,41^ While each of these single biomarkers provide predictive value, more complex models which capture nonlinear trends and multivariate relationships between biomarkers can provide improved predictive value across a variety of pathologies.

### Dyspnea is accurately predicted using machine learning

We show that dyspnea can be accurately identified using machine learning models and that this approach may produce more accurate estimations of self-reported dyspnea than human estimates via visual observation. Beyond the benefit of high recall and thus reducing false negative estimates, this model can provide continuously monitoring and notify healthcare providers when patients require intervention, particularly in-between routine check-ins when self-reported dyspnea can be obtained. In recent years, several groups have found that other subjective sensations such as pain can be predicted using a variety of input variables.^42–44^ Our findings align with this prior work, further suggesting that subjective sensations can be monitored or predicted using quantifiable inputs. These predictions are robust even given their relationship with psychological factors such as anxiety. We hypothesize that this is because anxiety modulates the input biomarkers in a predictable way and that underlying effects of anxiety on dyspnea are therefore captured.^45,46^ Robust, reliable prediction of subjective sensations such as pain and dyspnea across a variety of patients and pathologies can improve patient outcomes and reduce suffering, particularly in patients with communication barriers.

### Healthcare professionals under-estimate dyspnea severity

Despite the variety of self-reported cut-offs and RDOS thresholds tested, performance metrics for visual estimates of dyspnea were less accurate than our predictive model at distinguishing significant dyspnea. Precision performance between healthcare providers and our model were remarkably similar. However, the healthcare provider observational estimate recall was consistently lower. This indicates great certainty in the positive results from using the RDOS, but a general underestimation of patient dyspnea, and a large proportion of false negative estimates. In these cases, patients with less severe dyspnea, and those who show less obvious signs, may be left with unmanaged dyspnea. Lowering the threshold of the RDOS from scores of three to two dramatically increases recall (**Table S3**) with minimal loss of precision, leading to F1 and accuracy increases. This indicates that RDOS may be more beneficial for detecting any presence of dyspnea rather than a specific threshold to indicate moderate dyspnea or other varied intensities. This supports recent work by Aikawa et al. (2024) who demonstrate that the RDOS, IC-RDOS, and MV-RDOS can predict the presence of dyspnea with moderate accuracy, but is limited in its ability to discriminate severity, although work by Zhuang et al. (2018) show the RDOS can perform fairly well in discriminating moderate to severe dyspnea in palliative care patients.^32,47^ Aikawa et al. also demonstrated that lowering the cut off point for the RDOS was required to increase its performance, supporting our findings.

### Limitations

It is important to note that our current model was trained and tested with data from young participants without significant cardiovascular or pulmonary disease. Our dataset also includes only hypercapnia and hypoxia-induced dyspnea stimuli. Therefore, further studies must be done to include data across additional dimensions of dyspnea such as fixed ventilation patterns, airflow resistance, and check wall tightness. There may also be time-dependent impacts as our dataset includes 4-minutes of dyspnea stimuli so the effects of inspiratory muscle fatigue are not considered. Further studies should also verify the application of predictive multi-biomarker models for diverse patient populations exhibiting heightened airway resistance, chest wall restrictions, and other limitations and medical interventions which may modulate dyspnea severity (e.g. sedation and neuromuscular blockade).

One other limitation of our study was the inability to score nasal flaring in video recordings due to the use of opaque oronasal face masks. While the probability that this feature alone would have significantly impacted performance of observational estimates, and many adaptations of the RDOS exclude nasal flaring as a grading criterion, our findings should be interpreted in the context of this limitation.

## Conclusion

Machine learning is an excellent solution for the monitoring of subjective symptoms which manifest in measurable physiological biomarkers. Predictive models such as ours can be extracted and used as a lightweight addition to existing monitoring software or as standalone with a small, low power computer. Machine learning methods allow for the use of many variables to be considered simultaneously in a way that is difficult or impossible to do for unassisted humans. The application of machine learning to monitoring technologies such as those for identifying dyspnea also allows continuous monitoring between in-person patient visits. Future work will prioritize extending the current predictive model to include additional dimensions of dyspnea and validation in diverse clinical populations.

## Supporting information

All supplemental tables and figures

## Acknowledgements

We thank all study participants and healthcare providers for their time. We also thank members of the Heinrich lab who assisted with this study. A special thank you to Dr. Robert Banzett for his guidance.

## REFERENCES

1. von Leupoldt A, Sommer T, Kegat S, et al. The Unpleasantness of Perceived Dyspnea Is Processed in the Anterior Insula and Amygdala. Am J Respir Crit Care Med. 2008;177(9):1026–1032. doi:10.1164/rccm.200712-1821OC

2. Simon PM, Schwartzstein RM, Weiss JW, Fencl V, Teghtsoonian M, Weinberger SE. Distinguishable Types of Dyspnea in Patients with Shortness of Breath. Am Rev Respir Dis. 1990;142(5):1009–1014. doi:10.1164/ajrccm/142.5.1009

3. Schwartzstein RM, Manning HL, Weiss JW, Weinberger SE. Dyspnea: A sensory experience. Lung. 1990;168(1):185–199. doi:10.1007/BF02719692

4. Schmidt M, Banzett RB, Raux M, et al. Unrecognized suffering in the ICU: Addressing dyspnea in mechanically ventilated patients. Intensive Care Med. 2013;40(1):1. doi:10.1007/s00134-013-3117-3

5. Pesola GR, Ahsan H. Dyspnea as an Independent Predictor of Mortality. Clin Respir J. 2016;10(2):142–152. doi:10.1111/crj.12191

6. Parshall MB, Schwartzstein RM, Adams L, et al. An Official American Thoracic Society Statement: Update on the Mechanisms, Assessment, and Management of Dyspnea. Am J Respir Crit Care Med. 2012;185(4):435–452. doi:10.1164/rccm.201111-2042ST

7. Berliner D, Schneider N, Welte T, Bauersachs J. The Differential Diagnosis of Dyspnea. Dtsch Ärztebl Int. 2016;113(49):834–845. doi:10.3238/arztebl.2016.0834

8. Dyspnea. Am J Respir Crit Care Med. 1999;159(1):321–340. doi:10.1164/ajrccm.159.1.ats898

9. Dupuis-Lozeron E, Soccal PM, Janssens JP, Similowski T, Adler D. Severe Dyspnea Is an Independent Predictor of Readmission or Death in COPD Patients Surviving Acute Hypercapnic Respiratory Failure in the ICU. Front Med. 2018;5:163. doi:10.3389/fmed.2018.00163

10. Cammarota G, Simonte R, De Robertis E. Comfort During Non-invasive Ventilation. Front Med. 2022;9:874250. doi:10.3389/fmed.2022.874250

11. Worsham CM, Banzett RB, Schwartzstein RM. Dyspnea, Acute Respiratory Failure, Psychological Trauma, and Post-ICU Mental Health. Chest. 2021;159(2):749–756. doi:10.1016/j.chest.2020.09.251

12. von Leupoldt A, Dahme B. Psychological aspects in the perception of dyspnea in obstructive pulmonary diseases. Respir Med. 2007;101(3):411–422. doi:10.1016/j.rmed.2006.06.011

13. Frese T, Sobeck C, Herrmann K, Sandholzer H. Dyspnea as the Reason for Encounter in General Practice. J Clin Med Res. 2011;3(5):239–246. doi:10.4021/jocmr642w

14. Hayen A, Herigstad M, Pattinson KTS. Understanding dyspnea as a complex individual experience. Maturitas. 2013;76(1):45–50. doi:10.1016/j.maturitas.2013.06.005

15. Hui D, Morgado M, Vidal M, et al. Dyspnea in Hospitalized Advanced Cancer Patients: Subjective and Physiologic Correlates. J Palliat Med. 2013;16(3):274–280. doi:10.1089/jpm.2012.0364

16. Fukushi I, Pokorski M, Okada Y. Mechanisms underlying the sensation of dyspnea. Respir Investig. 2021;59(1):66–80. doi:10.1016/j.resinv.2020.10.007

17. Burki NK, Lee LY. Mechanisms of Dyspnea. Chest. 2010;138(5):1196–1201. doi:10.1378/chest.10-0534

18. Gigliotti F. Mechanisms of dyspnea in healthy subjects. Multidiscip Respir Med. 2010;5(3):195–201. doi:10.1186/2049-6958-5-3-195

19. Adams L, Lane R, Shea SA, Cockcroft A, Guz A. Breathlessness during different forms of ventilatory stimulation: a study of mechanisms in normal subjects and respiratory patients. Clin Sci Lond Engl 1979. 1985;69(6):663–672. doi:10.1042/cs0690663

20. Manning HL, Schwartzstein RM. Pathophysiology of Dyspnea. N Engl J Med. 1995;333(23):1547–1553. doi:10.1056/NEJM199512073332307

21. Campbell ML, Templin T, Walch J. A Respiratory Distress Observation Scale for Patients Unable To Self-Report Dyspnea. J Palliat Med. 2010;13(3):285–290. doi:10.1089/jpm.2009.0229

22. Haugdahl HS, Storli SL, Meland B, Dybwik K, Romild U, Klepstad P. Underestimation of Patient Breathlessness by Nurses and Physicians during a Spontaneous Breathing Trial. Am J Respir Crit Care Med. 2015;192(12):1440–1448. doi:10.1164/rccm.201503-0419OC

23. Banzett RB, Schwartzstein RM. Dyspnea: Don’t Just Look, Ask! Am J Respir Crit Care Med. 2015;192(12):1404–1406. doi:10.1164/rccm.201508-1637ED

24. Baker KM, DeSanto-Madeya S, Banzett RB. Routine dyspnea assessment and documentation: Nurses’ experience yields wide acceptance. BMC Nurs. 2017;16:3. doi:10.1186/s12912-016-0196-9

25. Demoule A, Decavele M, Antonelli M, et al. Dyspnoea in acutely ill mechanically ventilated adult patients: an ERS/ESICM statement. Intensive Care Med. 2024;50(2):159–180. doi:10.1007/s00134-023-07246-x

26. Mkrtchyan K, Lohn B, Qazi A, Ma S, Heinrich E. Addressing Disparities Between Physician Rated and Self-Reported Dyspnea Using Machine Learning. Physiology. 2024;39(S1):2192. doi:10.1152/physiol.2024.39.S1.2192

27. Mkrtchyan K g., Lohn B, Qazi A, Heinrich E, Ma S. Non-invasive Prediction of Dyspnea Severity Using Machine Learning. In: B80-2. ABNORMALITIES IN STRUCTURE AND FUNCTION IN COPD. American Thoracic Society International Conference Abstracts. American Thoracic Society; 2024:A4545–A4545. doi:10.1164/ajrccm-conference.2024.209.1_MeetingAbstracts.A4545

28. Heinrich EC, Orr JE, Gilbertson D, et al. Relationships Between Chemoreflex Responses, Sleep Quality, and Hematocrit in Andean Men and Women. Front Physiol. 2020;11. doi:10.3389/fphys.2020.00437

29. Palliative Care Benchmarks from Academic Medical Centers. doi:10.1089/jpm.2006.0048

30. Wysham NG, Miriovsky BJ, Currow DC, et al. Practical Dyspnea Assessment: Relationship Between the 0–10 Numerical Rating Scale and the Four-Level Categorical Verbal Descriptor Scale of Dyspnea Intensity. J Pain Symptom Manage. 2015;50(4):480–487. doi:10.1016/j.jpainsymman.2015.04.015

31. Campbell ML, Templin TN. Intensity cut-points for the Respiratory Distress Observation Scale. Palliat Med. 2015;29(5):436–442. doi:10.1177/0269216314564238

32. Aikawa G, Imanaka R, Sakuramoto H, Hatozaki C, Unoki T, Okamoto S. Assessment of Dyspnea in Critically Ill Patients: A Comparative Analysis of Evaluation Scales. Cureus. 16(1):e52751. doi:10.7759/cureus.52751

33. Wong RX, Shirlynn H, Koh YS, Goh Seow Lin S, Quah D, Zhuang Q. Exploration and Development of a Simpler Respiratory Distress Observation Scale (modRDOS-4) as a Dyspnea Screening Tool: A Prospective Bedside Study. Palliat Med Rep. 2021;2(1):9–14. doi:10.1089/pmr.2020.0094

34. Decavèle M, Rozenberg E, Niérat MC, et al. Respiratory distress observation scales to predict weaning outcome. Crit Care. 2022;26:162. doi:10.1186/s13054-022-04028-7

35. Moosavi SH, Golestanian E, Binks AP, Lansing RW, Brown R, Banzett RB. Hypoxic and hypercapnic drives to breathe generate equivalent levels of air hunger in humans. J Appl Physiol. 2003;94(1):141–154. doi:10.1152/japplphysiol.00594.2002

36. Buchanan GF, Richerson GB. Role of chemoreceptors in mediating dyspnea. Respir Physiol Neurobiol. 2009;167(1):9–19. doi:10.1016/j.resp.2008.12.002

37. Banzett RB, Lansing RW, Evans KC, Shea SA. Stimulus-response characteristics of CO2-induced air hunger in normal subjects. Respir Physiol. 1996;103(1):19–31. doi:10.1016/0034-5687(95)00050-X

38. Campbell ML. Psychometric Testing of a Respiratory Distress Observation Scale. J Palliat Med. 2008;11(1):44–50. doi:10.1089/jpm.2007.0090

39. Izumizaki M, Masaoka Y, Homma I. Coupling of dyspnea perception and tachypneic breathing during hypercapnia. Respir Physiol Neurobiol. 2011;179(2-3):276–286. doi:10.1016/j.resp.2011.09.007

40. Manning HL, Shea SA, Schwartzstein RM, Lansing RW, Brown R, Banzett RB. Reduced tidal volume increases ‘air hunger’ at fixed PCO2 in ventilated quadriplegics. Respir Physiol. 1992;90(1):19–30. doi:10.1016/0034-5687(92)90131-f

41. Banzett RB, Lansing RW, Binks AP. Air Hunger: A Primal Sensation and a Primary Element of Dyspnea. Compr Physiol. 2021;11(2):1449–1483. doi:10.1002/j.2040-4603.2021.tb00156.x

42. Hsiao FJ, Chen WT, Pan LLH, et al. Machine learning–based prediction of heat pain sensitivity by using resting-state EEG. Front Biosci-Landmark. 2021;26(12):1537–1547. doi:10.52586/5047

43. Lee J, Mawla I, Kim J, et al. Machine learning-based prediction of clinical pain using multimodal neuroimaging and autonomic metrics. Pain. 2019;160(3):550–560. doi:10.1097/j.pain.0000000000001417

44. Lötsch J, Ultsch A. Machine learning in pain research. Pain. 2018;159(4):623–630. doi:10.1097/j.pain.0000000000001118

45. Ritsert F, Elgendi M, Galli V, Menon C. Heart and Breathing Rate Variations as Biomarkers for Anxiety Detection. Bioengineering. 2022;9(11):711. doi:10.3390/bioengineering9110711

46. Tipton MJ, Harper A, Paton JFR, Costello JT. The human ventilatory response to stress: rate or depth? J Physiol. 2017;595(17):5729–5752. doi:10.1113/JP274596

47. Zhuang Q, Yang GM, Neo SHS, Cheung YB. Validity, Reliability, and Diagnostic Accuracy of the Respiratory Distress Observation Scale for Assessment of Dyspnea in Adult Palliative Care Patients. J Pain Symptom Manage. 2019;57(2):304–310. doi:10.1016/j.jpainsymman.2018.10.506

