## Supplementary material for "A Machine Learning Approach to Predicting Dyspnea with Noninvasive Biomarkers": All supplemental tables and figures

| $P_{IO2}$ (mmHg) | $P_{ETCO2}$ (mmHg) | Self-reported Dyspnea Severity | | |
|---|---|---|---|---|
| | | **Mean ± SD** | **Minimum** | **Maximum** |
| 149 | 45 | 0.961 ± 1.320 | 0 | 6.136 |
| 149 | 52 | 2.708 ± 2.181 | 0 | 8.887 |
| 80 | 45 | 1.763 ± 1.909 | 0 | 6.121 |
| 80 | 52 | 3.791 ± 2.461 | 0 | 8.5376 |
| 68 | 45 | 2.651 ± 2.186 | 0 | 8.010 |
| 68 | 52 | 4.542 ± 2.109 | 0.158 | 9.187 |

**Table S1.** Variation in dyspnea severity across the participant pool for each treatment group.

| | AUC Values | | Values Based on Ideal Threshold | | | |
|---|---|---|---|---|---|---|
| **Cut-Off** | **ROC** | **PR** | **Precision** | **Recall** | **F1** | **Threshold** |
| **2** | 0.882 | 0.783 | 0.776 | 0.804 | 0.789 | 0.35 |
| **2.5** | 0.865 | 0.726 | 0.644 | 0.826 | 0.724 | 0.33 |
| **3** | 0.923 | 0.832 | 0.816 | 0.795 | 0.805 | 0.42 |
| **3.5** | 0.864 | 0.666 | 0.535 | 0.697 | 0.605 | 0.36 |
| **4** | 0.888 | 0.649 | 0.652 | 0.556 | 0.600 | 0.51 |
| **4.5** | 0.928 | 0.601 | 0.475 | 0.905 | 0.623 | 0.25 |
| **5** | 0.903 | 0.576 | 0.405 | 0.833 | 0.545 | 0.29 |

**Table S2.** Summary of random forest performance across a range of NRS scale self-reported dyspnea cut-offs. The ideal F1 score was found at a cut-off of 3 with a threshold of 0.42.

| Dyspnea Cut-Off | RDOS Threshold | True Pos. | True Neg. | False Pos. | False Neg. | Precision | Recall | F1 | Accuracy |
|---|---|---|---|---|---|---|---|---|---|
| 2 | 1 | 49 | 4 | 9 | 6 | 0.845 | 0.891 | 0.867 | 0.779 |
| 2 | 2 | 40 | 7 | 6 | 15 | 0.870 | 0.727 | 0.792 | 0.691 |
| 2 | 3 | 27 | 11 | 2 | 28 | 0.931 | 0.491 | 0.643 | 0.559 |
| 2.5 | 1 | 46 | 6 | 12 | 4 | 0.793 | 0.92 | 0.852 | 0.765 |
| 2.5 | 2 | 37 | 9 | 9 | 13 | 0.804 | 0.740 | 0.771 | 0.676 |
| 2.5 | 3 | 24 | 13 | 5 | 26 | 0.828 | 0.480 | 0.608 | 0.544 |
| 3 | 1 | 46 | 6 | 12 | 4 | 0.793 | 0.920 | 0.852 | 0.765 |
| 3 | 2 | 37 | 9 | 9 | 13 | 0.804 | 0.740 | 0.771 | 0.676 |
| 3 | 3 | 24 | 13 | 5 | 26 | 0.828 | 0.480 | 0.608 | 0.544 |
| 3.5 | 1 | 39 | 6 | 19 | 4 | 0.672 | 0.907 | 0.772 | 0.662 |
| 3.5 | 2 | 33 | 12 | 13 | 10 | 0.717 | 0.767 | 0.742 | 0.662 |
| 3.5 | 3 | 22 | 18 | 7 | 21 | 0.759 | 0.512 | 0.611 | 0.588 |
| 4 | 1 | 39 | 6 | 19 | 4 | 0.672 | 0.907 | 0.772 | 0.662 |
| 4 | 2 | 33 | 12 | 13 | 10 | 0.717 | 0.767 | 0.742 | 0.662 |
| 4 | 3 | 22 | 18 | 7 | 21 | 0.759 | 0.512 | 0.611 | 0.588 |

**Table S3.** Enlargement of Table 2 showing physician performance with a variety of both self-reported dyspnea cut offs as determinates of significant dyspnea as well as RDOS scoring of significant dyspnea thresholds.

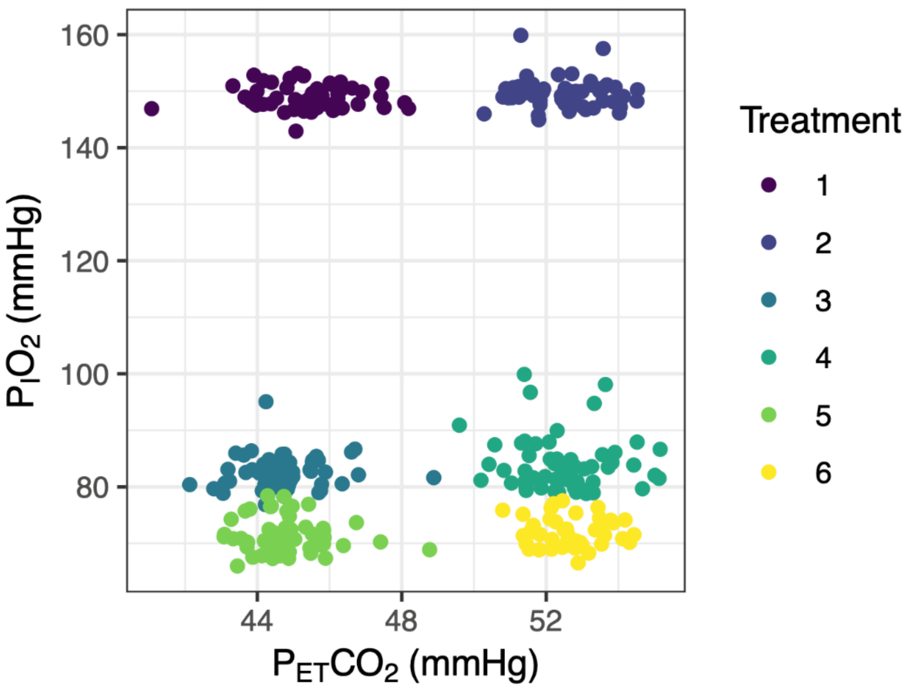


**Figure S1. Performance of end-tidal forcing experiments.** Individual data points represent the actual $P_{ET}CO_2$ and $P^{I}O_2$ values which each participant stabilized at during the last minute of the treatment period. Data point colors represent the target treatment: Treatment 1: $P_{I}O_2$ = 149 mmHg, $P_{ET}CO_2$ = 45 mmHg; Treatment 2: $P_{I}O_2$ = 149 mmHg, $P_{ET}CO_2$ = 52 mmHg; Treatment 3: $P_{I}O_2$ = 80 mmHg, $P_{ET}CO_2$ = 45 mmHg; Treatment 4: $P_{I}O_2$ = 80 mmHg, $P_{ET}CO_2$ = 52 mmHg; Treatment 5: $P_{I}O_2$ = 68 mmHg, $P_{ET}CO_2$ = 45 mmHg; Treatment 6: $P_{I}O_2$ = 68 mmHg, $P_{ET}CO_2$ = 52 mmHg.

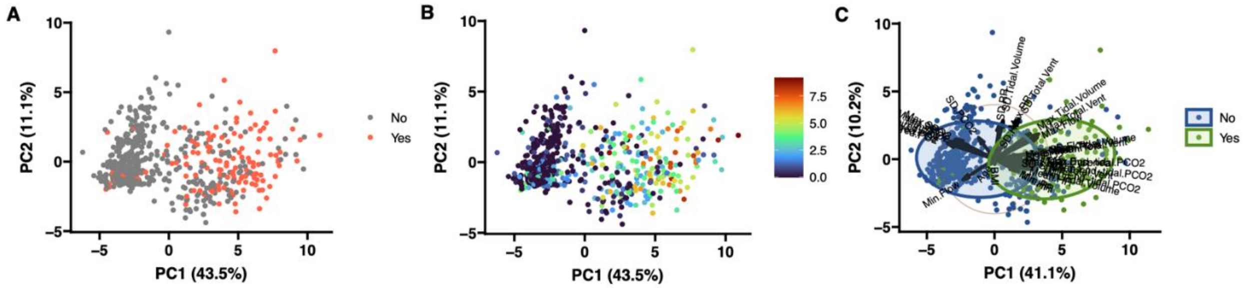


**Figure S2. Principal component analysis (PCA) plots.** Plots demonstrate separation of groups with severe dyspnea (self-reported NRS ≥3) and no significant dyspnea based on biomarker input data and basic demographic information (A). (B) Self-reported dyspnea scores colored as a continuous variable. (C) Ellipses highlighting separation of groups and vectors showing the impact of individual variables on the separation of groups.
